# PhyloMatcher: a tool for resolving conflicts in taxonomic nomenclature

**DOI:** 10.1101/2023.08.07.552263

**Authors:** Jonathan A. Rader, Madelyn A. Pivovarnik, Matias E. Vantilburg, Logan S. Whitehouse

## Abstract

**Summary:** Large-scale comparative studies rely on the application of both phylogenetic trees and phenotypic data, both of which come from a variety of sources, but due to the changing nature of phylogenetic classification over time, many taxon names in comparative datasets do not match the nomenclature in phylogenetic trees. Manual curation of taxonomic synonyms in large comparative datasets can be daunting. To address this issue, we introduce PhyloMatcher, a tool which allows for programmatic querying of two commonly used taxonomic databases to find associated synonyms with given target species names.

**Availability and implementation:** PhyloMatcher is easily installed as a Python package with pip, or as a standalone GUI application. PhyloMatcher source code and documentation are freely available at https://github.com/Lswhiteh/PhyloMatcher, the GUI application can be downloaded from the Releases page.

**Supplemental Information:** We provide documentation for PhyloMatcher, including walkthrough instructions for the GUI application on the Releases page of https://github.com/Lswhiteh/PhyloMatcher.

## Introduction

Large datasets and broad comparative studies have become the research paradigm across multiple disciplines of biology (Pal *et al*., 2020; da Silva *et al*., 2019; Muñoz and Price, 2019; Xia *et al*., 2020; Stephens *et al*., 2015; Yu and Nielsen, 2019; Tolani *et al*., 2021; Wüest *et al*., 2020). The rise of -omics technologies and high-throughput data collection permit rapid collection of data reflecting a wide variety of physiological, morphological, behavioral, and ecological traits (e.g., Modahl *et al*., 2020; Li *et al*., 2020; Chen *et al*., 2021; Nilsson *et al*., 2019). Genomic methods and the decreasing cost of sequencing have also made it possible to assemble large-scale phylogenies that will increasingly facilitate comparative exploration of trait data to illuminate macroevolutionary and macroecological patterns (e.g., Kim *et al*., 2021; Jetz *et al*., 2012; Prum *et al*., 2015; Varga *et al*., 2019; Buckley *et al*., 2010; Feng *et al*., 2020; Suvorov, Kim, *et al*., 2022; Suvorov, Scornavacca, *et al*., 2022). Despite the newfound abundance of comparative datasets and phylogenies, data that originate from different sources via disparate techniques are not always immediately compatible, and require careful curation prior to subsequent analysis (Marx, 2013; Leonelli, 2019; Pal *et al*., 2020; Xia *et al*., 2020).

A key necessity for broad studies of macroecology and trait evolution is that comparative data for each taxon must be matched to the appropriate branch tips within phylogenetic trees, which is confounded by mismatches of names between the dataset and the tree (Figure 1).

**Figure 1.**
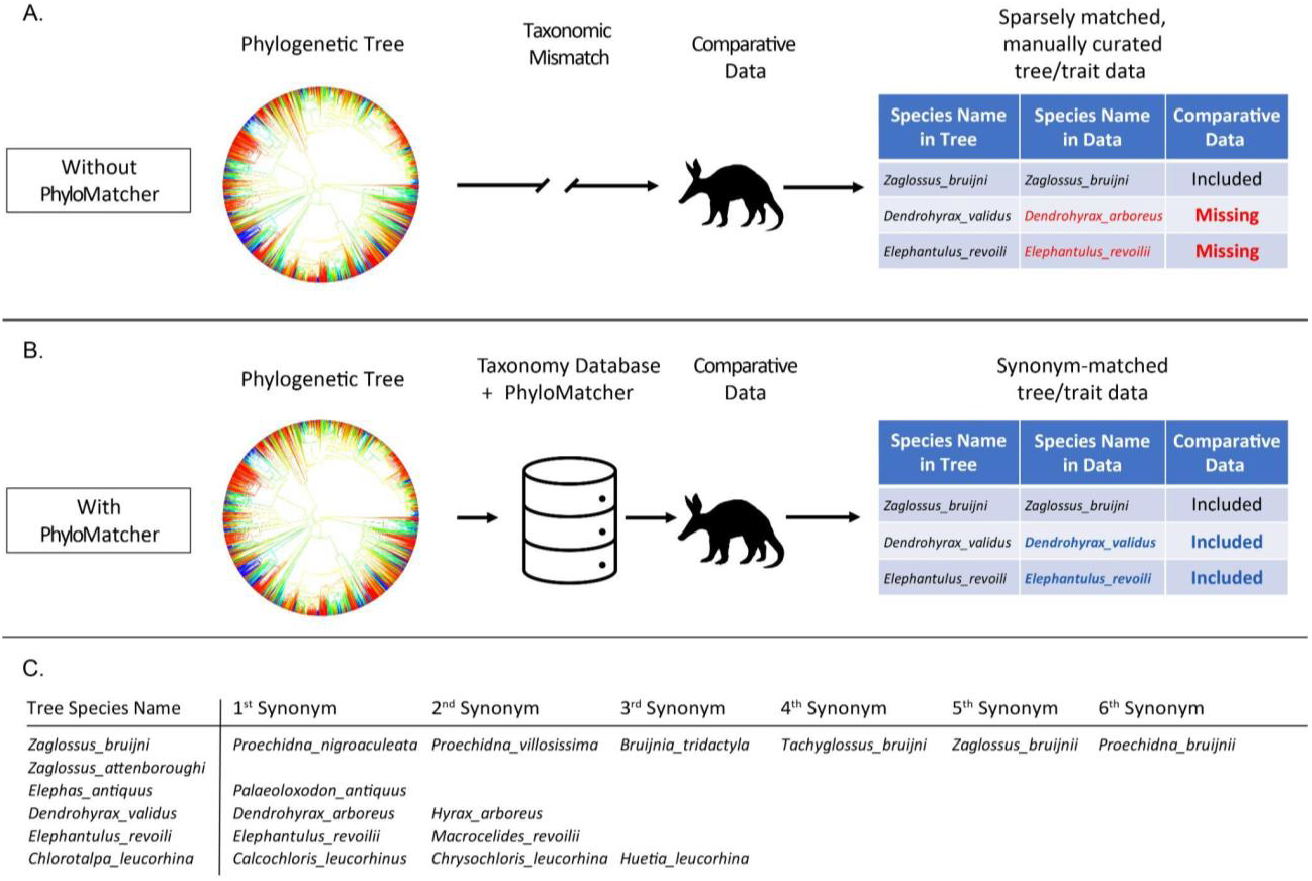
Mismatch of taxonomic nomenclature between species tips in phylogenetic trees and comparative datasets can lead to missing data and taxa being dropped from comparative analysis (A). We present a programmatic tool, PhyloMatcher, that can be used as an intermediary step in comparative workflows (B), to reconcile taxonomic mismatches, increasing the accuracy, completeness, and speed of comparative analyses. PhyloMatcher functions by compiling candidate synonyms for species names from a phylogenetic tree (C), and then reassigning names in the comparative dataset to match names in the tree.

Taxonomic naming conventions have fluctuated through time (e.g., Pauly *et al*., 2009; Eardley and Urban, 2006), and many species have a multitude of synonymous names (Patterson *et al*., 2010; Garnett *et al*., 2020). For example, research collections in natural history museums around the world provide a unique repository of biological data that has been accumulated for centuries. These collections are vast in both their taxonomic breadth and in the number of samples available for many taxa, facilitating broad comparative studies of a multitude of traits.

Furthermore, because specimens were collected at various times in history, they can provide opportunities for time series analyses that would not otherwise be possible (Holmes *et al*., 2016; Lister, 2011; Lopez *et al*., 2020; Shultz *et al*., 2021; though this is not without challenges, see Davis *et al*., 2023). Museum specimens have been labeled with these various synonyms through time - an artifact which has been inherited by comparative datasets compiled from the specimens. Similar problems can pervade field-collected datasets. The resulting taxonomic ambiguity poses a serious challenge to phylogenetic comparative analyses. The problem is easily rectified manually with small numbers of taxa, but it becomes less tractable as the number of study taxa increases. In especially taxonomically-broad studies, reconciling taxonomic names can impose a serious impediment.

Because of the pervasiveness of mismatched taxonomy throughout biological data, the ability to programmatically reconcile names as an initial step in comparative analyses would be valuable. There are approaches that use fuzzy-matching in Oracle PL/SQL to cure misspellings and typographic errors among taxonomic names (e.g., Rees, 2014), but to our knowledge, there are no freely-available utilities or services that match and reconcile synonymous taxonomic names. Repositories for naming synonyms already exist: the Global Biodiversity Information Facility (GBIF: The Global Biodiversity Information Facility, 2023) and National Center for Biotechnology Information Taxonomy (NCBI Taxonomy Project; Federhen, 2003) databases list known current and historical names that have been applied to each taxon. Thus, there are resources available to identify mismatched names, but their use still requires cumbersome manual searching and matching of synonyms to reconcile them with the naming convention used in any given phylogeny.

Here, we present a tool for programmatically matching taxonomic names between datasets: PhyloMatcher, which we have made available as a Python command line tool and a GUI application. We first describe the software’s algorithm (Figure 1) and then describe how we used PhyloMatcher to reconcile synonymous names for a dataset of mammalian taxonomy with the naming conventions from the phylogenies provided at VertLife.org (Jetz *et al*., 2012; Upham *et al*., 2019; Jetz and Pyron, 2018; Tonini *et al*., 2016; Stein *et al*., 2018). We provide a comparison between manual matching of names with the output of TaxonMatcher in both accuracy and speed of matching. We show that our method provides a dramatic reduction in the time needed to link species-level data to phylogenetic trees for comparative analyses, thus overcoming a significant barrier to the analysis of large comparative datasets.

## Methods

The initial implementation of PhyloMatcher draws from two sources (GBIF and NCBI) to populate lists of synonymous taxonomy (Figure 1). These two databases differ in their structure, so we used different approaches to extract names from each of them. We describe both implementations in the following sections, followed by a brief example of an application of PhyloMatcher.

### GBIF Implementation

PhyloMatcher uses the `species` module of the `pygbif` (GBIF: The Global Biodiversity Information Facility, 2023) package as an API client to query GBIF for synonyms. For a given species in the input file, the GBIF backbone database is first queried to retrieve the base-level identity for the target species using the `name_backbone` function. Upon successful retrieval the current name used by GBIF is stored and the unique species identifier is used to access any known synonyms using the `name_usage` function. If a “canonical” name is found that is different from the target name, it is returned along with the synonyms and current name. This process is done in parallel using the `concurrent.futures` package in Python with a user-specified number of threads, 4 being the default. Once all queries are complete, the results are collated into a CSV file where the first column is the input species name and each successive column is a synonym found in the GBIF database.

### NCBI Implementation

The NCBI querying module of `PhyloMatcher` uses the `Entrez` module (Schuler *et al*., 1996) of the `Biopython` package (Cock *et al*., 2009) as an API client to query the NCBI Taxonomy database for synonyms. Similar to the GBIF module, the database identifier for a given input species is retrieved using the `run_esearch` function. Due to the strictness of NCBI’s search, in the case of a failure during this step, a few common letter substitutions are searched (such as swapping “um” for “us” in *Orthriophis taeniurus*, for example). Once all possible identifiers have been retrieved, the list is checked for duplicate and non-valid entries, both of which are removed. XML entries for all filtered IDs are retrieved and parsed simultaneously using the `efetch` and `read` functions. For each entry in the resulting data the “OtherNames” and “Name” fields are exhaustively searched, and all unique entries are returned. Results are collated and output identically to the GBIF module explained above.

### Merging the Phylogeny with Comparative Data

Once synonyms have been retrieved a separate module allows for matching comparative data to the phylogenetic tips found in the original data. This module takes in two files: a csv file containing comparative values where the first column is the binomial species name (genus and species, separated by underscores), and the output csv produced by `PhyloMatcher`’s querying module. The remaining columns in the comparative file can be any data associated with each species such as trait data, occurrence observations, or a multitude of other species-level data to be used in comparative analyses. The synonym lists are stored as a list-of-lists (Fig. 1), and the comparative data are loaded line-by-line. For each line in the comparative file the listed species is checked against the species lists, and if present, is swapped for the name in the phylogeny. A modified version of the comparative data file is written containing a column with the phylogeny-matched species names.

### Programmatic vs. Manual Matching

To test the functionality and utility of PhyloMatcher, we matched taxonomic names for a dataset of approximately 24 million occurrence observations of mammals with 15,082 unique species names from the GBIF database with taxonomic names from the mammalian phylogeny from VertLife.org both manually and using PhyloMatcher. The tree contains 5,911 of the approximately 6,000 extant mammal species (Upham *et al*., 2019), suggesting that there are a large number of taxonomic synonyms present in the occurrence data. We took two approaches to matching the taxonomy in the dataset to the tree, a standard manual approach and PhyloMatcher. In both approaches, we searched the NCBI and GBIF databases for species names in the GBIF mammal occurrence data that do not match tip names in the phylogeny from Vertlife.org (Upham *et al*., 2019). We tracked the time spent manually collecting synonymous names for the mammal dataset, which we compared to the run-time of PhyloMatcher given the same input. We further compared the number of candidate naming synonyms found by each method and the proportion of missing data that was recovered.

## Results

There were 947 out of 5,911 (16%) species in the mammalian phylogeny that were not matched by species occurrence observations in the GBIF dataset. Some species were simply not represented in the occurrence data, but others were the product of mismatched taxonomy.

Running PhyloMatcher produced 1,303 unique candidate naming synonyms, preserving data from 379 species that would have been dropped from subsequent analysis (40% of the missing species). We then replaced synonyms in the occurrence data (∼24 million data points) with names matching the tree, resulting in a reconciled occurrence dataset ready for comparative analysis. The name reconciliation in the occurrence dataset took roughly 600 seconds of compute time on a single core.

By contrast, our manual effort produced a total of 646 candidate naming synonyms, effectively reconciling 165 species that would have been otherwise dropped from comparative analyses because of mismatched nomenclature (17.4% of the total number of missing data), leading to a loss of 214 species compared to PhyloMatcher. Additionally, the manual reconciliation of names took 6.8 hours, compared to 107 seconds of computational time on 8 computing cores for PhyloMatcher for the query step. In summary, Phylomatcher not only found more candidate synonyms to match taxonomic names in the mammalian occurrence data to the phylogeny than manual matching found and preserved more than double the number of dropped taxa that manual matching was able to, it did so in a small fraction of the time.

## Discussion

Here we provide a solution to a longstanding challenge to large-scale comparative studies: the problem of mismatched taxonomy (Dayrat, 2005; Patterson *et al*., 2010; Garnett *et al*., 2020). Taxonomic conflicts have slowed advances in multiple fields of biology, including phylogeography, evolutionary biology and population genetics, comparative morphology and biomechanics, and ecology (Bortolus, 2008; Dayrat, 2005). Efforts to quantify biodiversity (McNeely, 2002; Schlick-Steiner *et al*., 2010) and the study of pathogens (especially fungi; Wu *et al*., 2019; Almeida-Silva *et al*., 2021) have been especially impacted by ambiguous taxonomy. We created PhyloMatcher to resolve many of these conflicts. The PhyloMatcher software is available both as a command line program and a GUI application, and provides a programmatic means to reconcile taxonomic names in trait data with the species labels in phylogenetic trees.

PhyloMatcher is easy to install and incorporate into existing workflows, and leverages pre-existing and continually maintained databases that include lists of synonymous species names. The input for PhyloMatcher is a list of taxonomic names from a phylogenetic tree and a trait dataset to be matched to it. The output is a new data file with reconciled species names, ready for comparative analyses. In our validation tests, PhyloMatcher roughly doubled the number of candidate matches in our test dataset relative to manual matching, and in a small fraction of the time. With the rise of large datasets and taxonomically-broad phylogenetic comparative analyses, tools that enhance the integration of multiple data types are increasingly important. Incorporating PhyloMatcher as a simple intermediate step in pre-existing workflows will improve the speed, accuracy, and phylogenetic completeness of comparative biology studies.

## Acknowledgements

We thank Daniel Matute, Dan Schrider, Evan Drage, Tyler Kent, Andrius Dagilis, Sean Anderson, and members of the Matute and Schrider labs for their helpful comments on this manuscript. Funding for this work was provided by the National Institutes of Health under award R01AI153523.

